# NeuralDock: Rapid and conformation-agnostic docking of small molecules

**DOI:** 10.1101/2021.08.20.457163

**Authors:** Congzhou M. Sha, Jian Wang, Nikolay V. Dokholyan

## Abstract

Virtual screening is a cost- and time-effective alternative to traditional high-throughput screening in the drug discovery process. Both virtual screening approaches, structure-based molecular docking and ligand-based cheminformatics, suffer from computational cost, low accuracy, and/or reliance on prior knowledge of a ligand that binds to a given target. Here, we propose a neural network framework, NeuralDock which accelerates the process of high-quality computational docking by a factor of 10^6^, and does not require prior knowledge of a ligand that binds to a given target. By approximating both protein-small molecule conformational sampling and energy-based scoring, NeuralDock accurately predicts the binding energy and affinity of a protein-small molecule pair, based on protein pocket 3D structure and small molecule topology. We use NeuralDock and 25 GPUs to dock 937 million molecules from the ZINC database against superoxide dismutase-1 in 21 hours, which we validate with physical docking using MedusaDock. Due to its speed and accuracy, NeuralDock may be useful in brute-force virtual screening of massive chemical libraries and training of generative drug models.

## INTRODUCTION

Drug discovery as carried out by pharmaceutical companies requires an investment of years of research effort and billions of dollars^1^. The preclinical pipeline for identifying a small molecule ligand for a protein target is: (1) biochemical screening of small molecules against a protein target or cellular assay, (2) medicinal chemistry optimization of candidate small molecules, and (3) validation of promising molecules in animals^2^. Step (1) is critical in identifying small molecules which bind tightly to the target (leads) and for their subsequent optimization in step (2) (hits). Step (1) is expensive and time-consuming, taking several months to screen a small library of >10^5^ compounds. Insufficient binding affinities of leads and hits from steps (1) and (2) often lead to drug attrition in step (3) and subsequent clinical trials, with attrition rates as high as 95%^4,5^. Molecular dynamics and rational drug design can explore a larger part of the chemical space and potentially increase binding affinity of hits, but typical docking tools still only dock one compound every few minutes at moderate sampling accuracy (AutoDock Vina: 1.2 minutes^6^, DOCK 6: 4.8 minutes^7^, Glide 1.7 minutes^8^, MedusaDock: seconds to minutes^9^), while the chemical space of potential drugs may be as large as 10^60^ small molecules^10^. Neural networks have shown significant promise in structural biology, accurately reproducing gold standard results in a fraction of the time^11^. Here, we accelerate the virtual docking process by 6 orders of magnitude, enabling docking of 10^9^ compounds in a single day at low cost.

Modern methods of computational drug docking are implemented by tools, such as MedusaDock^12,13^, AutoDock Vina^6,14,15^, DOCK^7^, and Glide^16^. These tools perform molecular docking using classical force fields to evaluate the binding energy or affinity of a small molecule to a protein pocket of interest. Here, we focus on MedusaDock because it performs fully flexible conformational sampling of both the protein and ligand, which mimics the induced fit model of protein-small molecule binding, whereas other tools generate ensembles of the protein which are then rigidly fixed and docked to the small molecule (AutoDock Vina^17^, DOCK^7^, Glide^18^). MedusaDock consists of two independent tasks: conformational sampling and scoring. Here, we show that both tasks can be well-approximated by a deep neural network at a fraction of the computational cost of traditional docking. Although we used MedusaDock to generate our data, the framework we have developed can be applied to the results of other docking tools.

The principal advantages of a neural network over traditional docking tools include differentiability (propagation of gradients in model training through automatic differentiation) and speed. Neural networks are also valuable for their composability, in which they can be used as subnetworks of larger neural networks while providing gradients for training^19,20^. Neural network inference is highly optimized on modern processors, and particularly on GPUs. One can achieve many orders of magnitude higher performance with neural network approximation than with traditional algorithms based on exact calculation^21^. As molecular dynamics is already a significant approximation to a quantum mechanical and statistical reality, inaccuracies in neural network predictions may be acceptable for virtual docking purposes^22^.

Deep neural networks for predicting binding affinities have been successful, however there are drawbacks to specific approaches^23–26^. For example, Francoeur et al.^23^ proposed to approximate force fields using neural network-based scoring while still relying on extensive conformational sampling, hence retaining the major computational bottleneck of virtual docking. Gentile et al.^24^ used neural networks to aid in chemical screening, but the predictions were nonspecific: information about the protein pocket was not used in the screening^24^. K_DEEP_ by Jiménez et al.^25^ uses computationally expensive convolutional architectures which limit inference speed, and was trained directly on protein-small molecule 3D crystal structures. Due to the inclusion of protein-small molecule crystal structures, K_DEEP_ is biased and has limited generalizability to proteins not bound to small molecules. This issue of bias has been discussed in Francoeur et al.^23^ and arises from self-docking, in which the neural network is provided with a low conformational energy crystal structure as input, and therefore does not perform conformational sampling. Cang et al.^26^ use convolutional networks with manually constructed ligand features based on ligand topology, but provide no forward validation with docking tools; instead, only binding affinity is predicted, increasing the risk of overfitting and self-docking bias.

In stark contrast to work such as in Francoeur et al.^23^, we do not explicitly train our neural network to predict the energy of a specific conformation. Rather, we train our network in a conformation-invariant manner by withholding conformational information in the inputs and by predicting population parameters like minimum energy and binding affinity. Unlike in work from Jiménez et al.^25^ and Cang et al.^26^, we limit bias from self-docking and overfitting. We train the neural network to directly predict the minimum binding energy evaluated by MedusaDock, based on a coarse 3D representation of the protein and a graph representation of the small molecule. The direct prediction of binding energy makes conformational sampling implicit in the neural network. The coarse protein representation and graph representation of the small molecule withholds the optimal orientation, alignment, and conformation of the small molecule in the protein pocket of interest from the neural network. Since we augment our training data with MedusaDock energies, we also reduce the likelihood of overfitting. With these design elements in our neural network NeuralDock, we achieve class-leading performance, as tested on the PDBbind 2013 core set. We perform a proof-of-concept docking for benchmarking and external validation purposes, using NeuralDock to dock nearly 10^9^ molecules from the ZINC database against the enzyme superoxide dismutase-1 (SOD1), which is not present in the training, validation, or test sets. Finally, we validate the predicted energies using MedusaDock.

## RESULTS

### Validation of the ability of NeuralDock to predict MedusaDock energy and experimental pK

NeuralDock training on MedusaDock energies and experimental pKs converged, achieving agreement with experimental binding affinities for the test set, the PDBbind 2013 core set (Figure 1b), and the validation set taken from PDBbind 2017 (Figure 1c). As discussed by Francoeur et al.^23^, testing on the core set may offer a biased evaluation of binding affinity prediction performance. We provide the core set correlation for comparison with other methods which solely report that data (Table 1). Even with a relatively small training set (2331 structures) and the massive diversity of potential protein pockets and ligands, NeuralDock was able to learn the binding affinities so accurately. We believe that the key to NeuralDock’s success was using high quality data produced by MedusaDock, which sampled thousands of conformations for each protein-ligand pair, and using only coarse 3D protein information and the small molecule’s topology as inputs to NeuralDock. By using only the topology of the small molecule as input, we forced NeuralDock to approximate the effects of conformational sampling.

**Figure 1:**
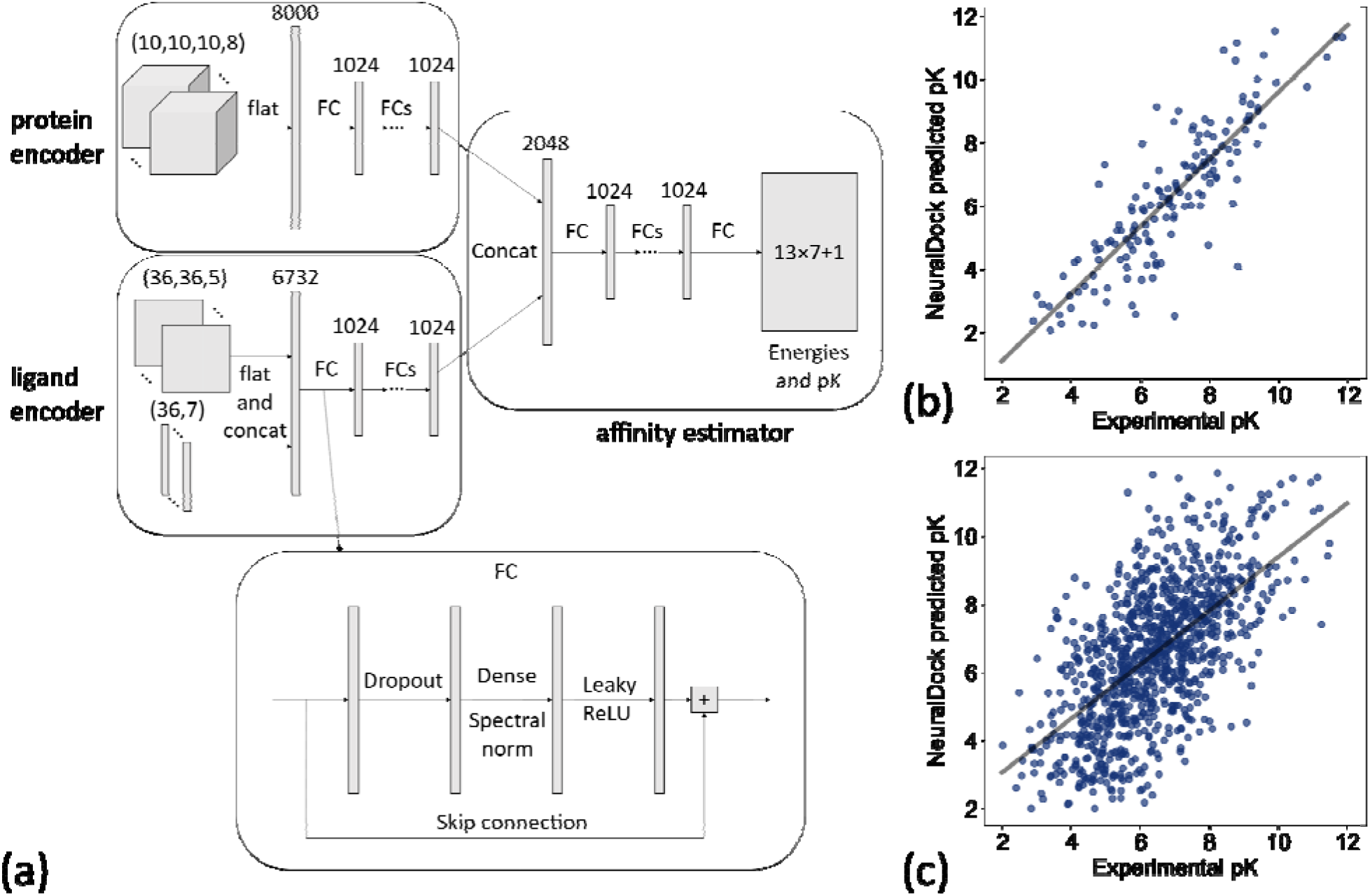
The neural network architecture on the left and performance comparison with MedusaDock on the right. (a) Inputs, hidden layers, and outputs are shown for the architecture. The protein pocket is flattened and fed into a subnetwork, and the ligand is processed similarly. The outputs of the two subnetworks are concatenated and fed into another subnetwork, which outputs 13×7+1 values representing the 7 summary statistics of the 13 energies output by MedusaDock, as well as the pK of the protein-ligand pair. The structure of each FC layer is shown at the bottom. (b) The 45 million parameter NeuralDock network achieves class-leading performance on the PDBbind 2013 core set (r=0.85, p<0.0001). (c) The 45 million parameter NeuralDock network achieves good agreement with experimentally determined pK on the validation set (r=0.62, p<0.0001).

**Table 1:** Correlation coefficients for binding affinity prediction of a variety of neural networks.

| Model | PDBbind core set<br>binding affinity<br>correlation | Number of test set<br>structures | Number of<br>training set<br>structures |
| --- | --- | --- | --- |
| Def2018 General<br>Ensemble<sup>23</sup> | 0.80 | 280 | 18,450 protein-ligand complexes<br>and 22,584,102<br>poses |
| K<sub>DEEP</sub><sup>25</sup> | 0.82 | 195 | 13,308 protein-ligand complexes |
| TopBP-ML<sup>26</sup> | **0.85** | 195 | 22,886<br>compounds<br>against each of<br>102 protein<br>targets |
| NeuralDock | **0.85** | 154 | 2,331 protein-ligand complexes |

To test the robustness, generalizability, and speed of NeuralDock, we performed virtual screening of a massive library of ligands (N=936,054,166) against the pocket at the dimeric interface of SOD1 and compared the results to MedusaDock (Figure 2). We compared relationships among MedusaDock E_without VDWR_, NeuralDock E_without VDWR_, and experimental binding affinities (pK), and found that they agree (Figure 3). E_without VDWR_ is the component of MedusaScore which is most highly correlated with experimental pK^27^. We demonstrate the correlation between NeuralDock and MedusaDock E_without VDWR_, first on the validation set drawn from PDBbind 2017 refined set (r=0.83, p<0.0001), and then on 100 random small molecules from ZINC docked to 1UXM, an A4V mutant of the human superoxide dismutase-1 enzyme (r=0.69, p<0.0001), (Figure 3a). Therefore, NeuralDock was successful in learning to predict the minimum E_without VDWR_ in a single shot. Additionally, we performed 2-way analysis of covariance (ANCOVA), which measures the effects of a categorical variable. In this instance, showed no statistically significant difference in the trends for the validation set and the external validation on 1UXM (F=0.67, p=0.41), which provides evidence that NeuralDock successfully generalizes from the training set to other protein-small molecule pairs, as well as achieving cross-docking success. Since the native structure 1UXM was not bound to any small molecule and MedusaDock performs fully flexible conformational sampling of both the protein and the small molecule, our results provide external validation of NeuralDock and demonstrate robustness under cross-docking, i.e. docking of small molecules to a protein as it appears in its native, unbound conformation.

**Figure 2:**
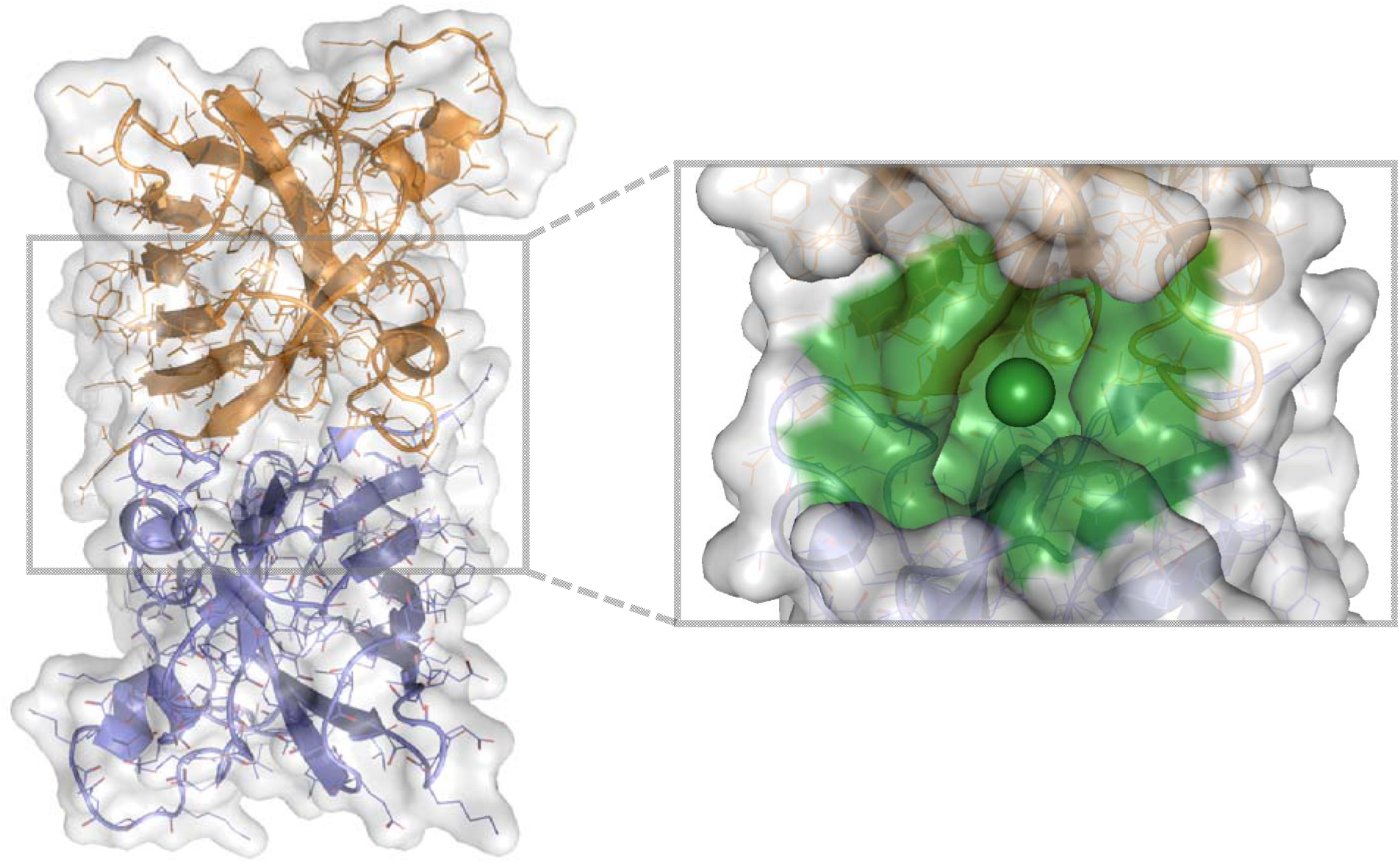
1UXM A4V SOD1 dimer chains A (gold) and B (lilac), with the protein pocket of interest (green) in billion-molecule docking. A cartoon, stick, and water-accessible surface representation of 1UXM, an A4V mutant SOD1 dimer structure. The image was generated using PyMOL 2.4.0^48^.

**Figure 3:**
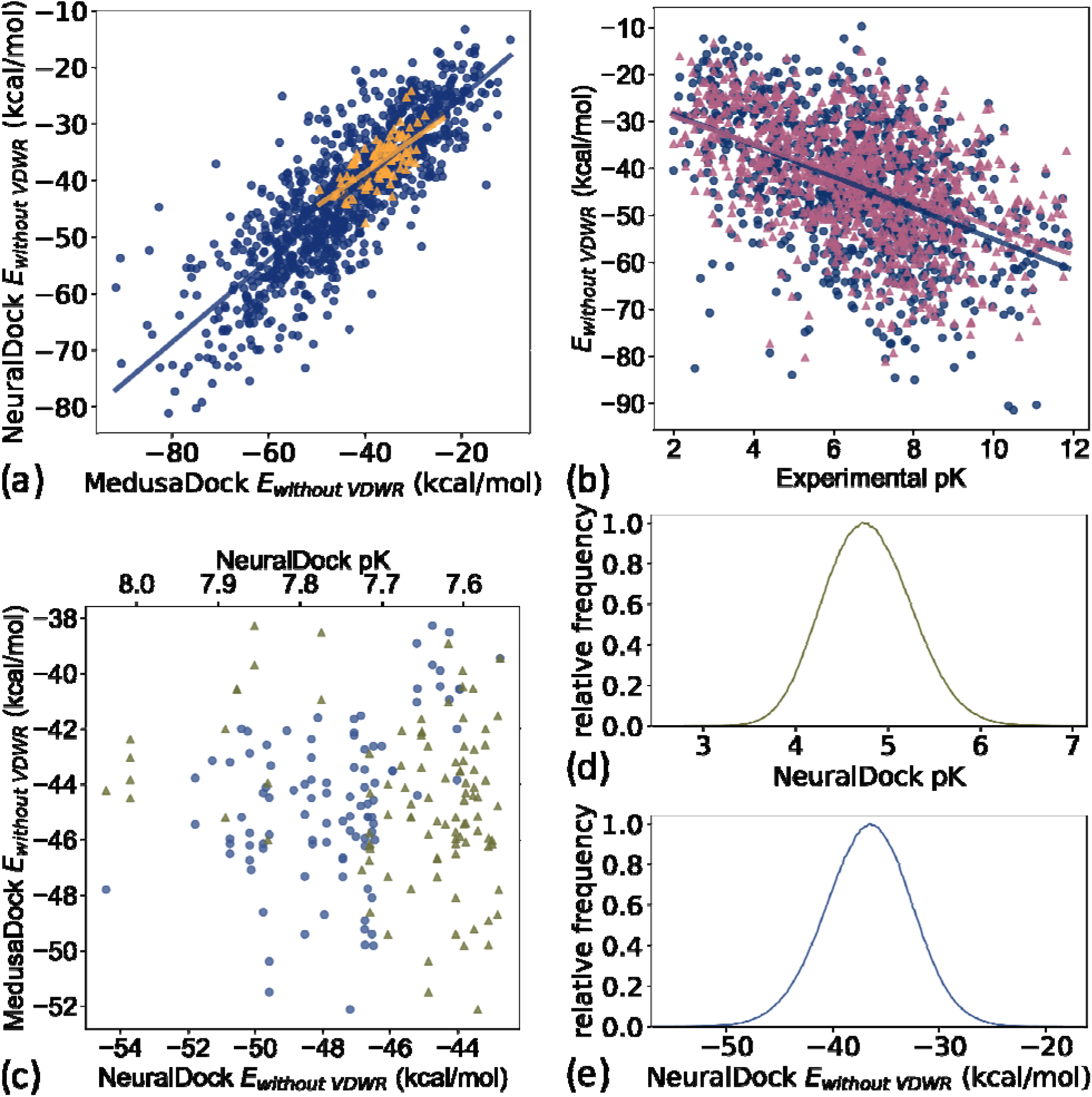
Comparisons among MedusaDock energies, NeuralDock predicted energies, and experimental binding affinity data. (a) The correlation between NeuralDock predicted E_without VDWR_ and MedusaDock E_without VDWR_ on the validation set (blue circles, r=0.83, p<0.0001) and 100 random small molecules docked to 1UXM (orange triangles, r=0.69, p<0.0001). NeuralDock performs well on the validation set in predicting MedusaDock energies, and the trend generalizes to 1UXM with no significant difference (2-way ANCOVA F=0.67, p=0.41). (b) The correlations of MedusaDock E_without VDWR_ (blue circles), NeuralDock predicted E_without VDWR_ (magenta triangles), and experimental binding affinity (pK) on the validation set (r=-0.48 for both data sets, p<0.0001), with no significant difference (2-way ANCOVA F=1.27, p=0.26). (c) The 100 small molecules with maximum NeuralDock pK (green triangles), from docking of 936,054,166 small molecules from the ZINC library against 1UXM; the corresponding predicted E_without VDWR_ is plotted (lilac circles). Left is higher binding affinity (higher pK) and lower energy (lower E_without VDWR_). (d) and (e) The relative frequency distributions (300 bins) of NeuralDock predicted pK (mean 4.07, std 0.47) and E_without VDWR_ (mean −36.6, std 4.1) respectively, on 8,099,176 (9% of total) randomly selected small molecules from the docking of 1UXM. The plots are centered at the means, the x-axis ranges are +/− 5 standard deviations from the mean, and colors are repeated from (c). Note that both the E_without VDWR_ and pKs in (c) are drawn from the extreme tails of the distributions shown in (d).

We found agreement of MedusaDock E_without VDWR_ and NeuralDock E_without VDWR_ with experimental pK, with similar correlations for both tools (r=-0.48 for both, p<0.0001), and no statistically significant difference in correlations with 2-way ANCOVA (F=1.27, p=0.26), (Figure 3b). The correlation of MedusaDock E_without VDWR_ with experimental pK (r=−0.48) is comparable to that of AutoDock Vina scoring (r=0.41) as reported in Francoeur et al.^23^, with MedusaDock performing better, likely due to our extensive sampling and computational effort for each protein-ligand pair. NeuralDock predicts experimental pK better than MedusaDock on the validation set (Figures 1b, 1c, 3b), however experimental validation of the pKs is needed to confirm that this result holds for SOD1 and other proteins. NeuralDock’s agreement with MedusaDock predictions demonstrates that it may be useful in replacing traditional docking tools such as MedusaDock and AutoDock Vina.

For the billion-molecule docking of 1UXM, we find that the 100 small molecules with the highest NeuralDock-predicted pKs also have low binding energies, with agreement of NeuralDock and MedusaDock E_without VDWR_ on those small molecules (Figure 3c). Furthermore, NeuralDock predicts that few other molecules have such high pK and low energy (Figures 3d). Combining these results with our validation set regression (Figure 1c), we predict that some of the 100 small molecules chosen have pK of approximately 8 (K≈10 nanomolar). Ninety-five of the 100 compounds satisfy Lipinski’s rule of five, and all have reasonable QED scores (median 0.50, range 0.33-0.66). The druglikeness of these compounds can be explained as bias from the ZINC Tranches, in which only 14,744,513 (1.5% of the over 997 million) compounds have molecular weight exceeding 500 Daltons or log P exceeding 5.

### Comparison of various NeuralDock architectures

Various architectures for NeuralDock yielded similar results (Table 2). We optimized the NeuralDock architecture based on E_without VDWR_ correlation, as E_without VDWR_ is most highly correlated with binding affinity^27^. Optimizing the network based on E_without VDWR_ prevents overfitting the hyperparameters (i.e. number of layers and total number of trainable parameters), as MedusaDock E_without VDWR_ is an external source of training information. Including other MedusaDock energies in training also helped to prevent overfitting (Supplementary Table 2). Even a limited network such as the 4.9 million parameter model was able to generalize from the training set to the validation set (Table 2).

**Table 2:** Correlation coefficients of NeuralDock predicted minimum energy and MedusaDock output for a variety of architectures. *We chose the 45 million parameter network as the best architecture; however this may change with larger and more diverse training sets.

| **Number of hidden**<br>**layers per subnetwork**<br>**(total)** | **Dimension of**<br>**hidden layer** | **Number of**<br>**trainable**<br>**parameters** | **Correlation**<br>**coefficient for $E_{\text{without}}$**<br>**VDWR** |
| --- | --- | --- | --- |
| 10 FC blocks (30 total) | 2048 | 83,221,686 | 0.794 |
| 10 FC blocks (30 total) | 1024 | 45,618,267 | 0.838* |
| 6 FC blocks (18 total) | 512 | 12,055,131 | 0.800 |
| 6 FC blocks (18 total) | 256 | 4,913,499 | 0.758 |
| 6 inception blocks (18<br>total) | N/A | 55,937,979 | 0.775 |

### NeuralDock dramatically accelerates large-scale docking-based virtual drug screening

Benchmarking on a single Tesla T4 GPU was able to predict energies and pKs of 96000 protein-small molecule pairs in 203.8 seconds, or 2 milliseconds per protein-small molecule pair. Docking of the 937 million ZINC compounds with NeuralDock took 21 hours on 25 GPUs, which matches our benchmarking of 2 milliseconds per protein-small molecule pair per GPU. For 3875 structures (including those which exceeded our computational resources), it took 120 processors 4 weeks to perform extensive MedusaDock sampling (1000 iterations), or an average of 20 hours per structure per processor. Taking 10 hours, as a conservative minimum compute time for MedusaDock, NeuralDock performs 1.7×10^7^ times faster. NeuralDock performs 10^5^ times faster than DOCK 6^7^, and 100 times faster than K_DEEP_^25^. NeuralDock is also faster than the Def2018 General Ensemble by Francoeur et al.^23^, since we directly predict binding affinity in a single forward pass instead of requiring many conformational samples and forward passes through the network^23^.

## DISCUSSION

There are several potential improvements to NeuralDock. First, NeuralDock takes a coarse atomic image of the 20 Å protein pocket as input. Including inputs such as protein amino acid composition, secondary structure, and hydrophobicity may improve NeuralDock predictions. This potential change in inputs is supported by the fact that NeuralDock was able to learn local contributions to the energy well (i.e. E_VWDA_, Supplementary Tables 1 and 2), while having difficulty with global contributions (e.g. E_solv_, Supplementary Tables 1 and 2). Second, NeuralDock is trained on a relatively small training set, and we did not use any small molecules which are off-target, i.e. do not to bind to the given protein. However, our external validation with MedusaDock indicates that this potential source of bias was at least partially addressed by the variety of binding affinities present in our training set. This deficiency may be remedied by docking more targets with MedusaDock as a reference, such as the PDBbind general set, which incurs a significant computational cost in data preparation. Additionally, the PDBbind general set experimental pKs may be noisier than in the refined set. Third, the binding energies and pK are invariant under rotations of the protein pocket as well as under permutations in the atom order for the small molecule bond adjacency matrix and atom type vector. Newer neural network architectures which directly leverage these physical symmetries have been proposed^28,29^.

MolGAN is a generative adversarial approach to drug discovery, which does not take into account the protein pocket^30,31^. Since NeuralDock is end-to-end differentiable, one can augment MolGAN training with NeuralDock acting as a scoring function to create drugs targeting specific proteins. The addition of such guidance may improve the stability of MolGAN training. While automatically differentiable force fields are currently being developed for molecular dynamics^32^, the advantage of NeuralDock is it can immediately evaluate the quality of a generated small molecule, implicitly performing conformational sampling, whereas conformational sampling must still be done for a differentiable force field. NeuralDock does not predict the explicit minimum energy configuration for the protein-small molecule pair, however, once specific compounds are identified via NeuralDock, the candidates can be docked by MedusaDock or another docking software prior to experimental validation, as the computational cost of doing so is much less than of traditional docking for the entire compound library.

## CONCLUSION

NeuralDock is a robust neural network for predicting binding energies and affinities for protein-small molecule pairs. The key design elements which result in its class-leading performance include high quality scoring of thousands of protein-small molecule conformations, coarse 3D image representation of the protein pocket, and topological representation of the small molecule. Because NeuralDock is trained to output predicted energies and pK in a single shot, NeuralDock is faster than competing methods and can be used in billion-molecule virtual screening. NeuralDock has been validated using the fully flexible docking tool MedusaDock and is ready for experimental validation.

## METHODS

### Data

For NeuralDock inputs, we used crystal structures of 3875 known protein-ligand pairs from the PDBbind 2017 refined set^33^, which were shuffled into a training set (N=2712, 70% of structures) and a validation set (N=1163, 30% of structures). One of these structures was not processable by MedusaDock 2.0, and 127 structures exceeded our computational resources. A further 496 ligands were not processable by rdkit 2020.09.3^34^. This left a training set of N=2279 (84.0% of original) and a validation set of N=972 (83.6% of original). The core set of PDBbind 2013 was used as the test set (N=195); 154 of these were processable by rdkit. There is no overlap of proteins among the training, validation, and test sets.

We extracted the atoms in the protein which were within a cube of side length 20 angstroms, centered at the ligand. These protein pockets were encoded as 10×10×10, 2-angstrom resolution images with 8 channels corresponding to a one-hot encoding of (no atom, C, O, N, S, P, H, other). If multiple atoms were contained in the same 2-angstrom cube, we took the maximum of each channel of the one-hot encodings. Using rdkit, we encoded the ligand as a length 36 atom type vector with 7 channels (no atom, C, N, O, F, S, other) and a 36×36 bond adjacency matrix with 5 channels (no bond, single, double, triple, aromatic/conjugated). For ligands with greater than 36 heavy atoms (hydrogens were excluded), we removed atoms with the least bond order until we were left with 36 total heavy atoms, thus attempting to preserve the important topologies present in the ligand. These input dimensions were chosen so that a comparable number of parameters in NeuralDock would be devoted to processing the protein and the ligand each (Figure 1a), as well as to prevent the ligand representation from becoming too sparse. We also chose a limit of 36 heavy atoms since molecules with greater than 36 heavy atoms are likely to exceed the 500 Dalton cutoff for Lipinski’s rule of five.

For each protein-ligand pair, we ran MedusaDock for 24 hours or 1000 iterations (whichever came first) on a single core of an Intel Xeon E5-2680 v3 processor with 6 GB RAM. We collected summary statistics (mean, median, standard deviation, minimum, maximum, skew, and kurtosis) on the 13 interaction energies computed in MedusaDock’s force field, MedusaScore^27^. Five of these energies were zero for most or all structures (see Supplementary Information). An interaction energy is defined as the total energy of the protein-ligand complex *E_P-L_* minus the contributions from the protein *E_P_* and the ligand *E_L_* when they are isolated.

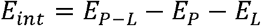

In developing our models, we chose to focus on E_without VDWR_, the interaction energy excluding repulsive van der Waals forces. E_without VDWR_ is the output of MedusaScore which is known to be most highly correlated with experimental binding affinity^27^.

The compounds (N = 997,402,117) were downloaded from the ZINC database^35^ (available in Tranches) and processed into the tensor input format described above. 936,054,166 (94%) of these compounds were processable by rdkit. We chose the 1UXM SOD1 PDB structure as a protein target^36^. The interface between the A and B chains (Figure 2) was chosen as the binding pocket for 1UXM.

### Neural network

For NeuralDock, we chose a fully-connected architecture, with spectral normalization^37^, dropout^38^ with rate 0.2, LeakyReLU activation^39^, and skip connections^40^. These characteristics were chosen as standard methods of model regularization to prevent overfitting and vanishing or exploding gradients during training.

The NeuralDock architecture (Figure 1a) was implemented in TensorFlow 2.4.0^41^ and Python 3.7. The inputs were the protein pocket and ligand topology, and the outputs were the 7 summary statistics of the 13 interaction energies computed by MedusaDock as well as the binding affinity pK=log_10_ K (dissociation/inhibitor constant K_D/I_ with units of molar). We considered K_D_ and K_I_ to be the same for the training purposes. We chose 10 fully-connected (FC) hidden layers for each of the three parts (protein encoder, ligand encoder, affinity predictor) of the network (Figure 1a), resulting in approximately 45 million trainable parameters. We varied the number of hidden layers as well as their widths during hyperparameter optimization (Table 2). The loss function was the L^2^ (squared difference) loss between the NeuralDock output energies and the MedusaDock output energies, as well as pKs. The Adam optimizer was used^42^, with a learning rate of 10^−6^. Training of each model took place on one NVIDIA Tesla T4 GPU, and the models were trained to convergence within a week. We trained a convolutional architecture, in which the FC blocks in the protein encoder (top left of Figure 1a) were replaced by spectrally normalized 3D inception modules^43^ with a comparable number of trainable parameters (Table 2).

### Chemical scoring

We used Lipinski’s rule of five^44^ to evaluate small molecule lead quality and druglikeness, as well as the Quantitative Estimate of Druglikeness (QED)^45^. Octanol-water partition coefficients (log P) were extracted from HTML files of the ZINC database, while all other quantities were computed using rdkit^34^.

### Statistics

Least squares regression was performed in Python 3.7 using SciPy 1.6.0^46^. Analysis of covariance (ANCOVA) was performed in Python 3.7 using Pingouin 0.3.12^47^.

## Supporting information

Supplementary Information for NeuralDock

## DATA AVAILABILITY

The data that supports the findings of this study are available from the corresponding author upon reasonable request.

## ACKNOWLEDGMENTS

We acknowledge support from the National Institutes for Health (R35 GM134864), National Science Foundation (2040667) and the Passan Foundation. We are thankful to the NSF Convergence Accelerator (award number 2040667) team for helpful discussions on machine learning. The content is solely the responsibility of the authors and does not necessarily represent the official views of the NIH.

## AUTHOR CONTRIBUTIONS

CMS implemented, trained, and validated the neural networks; performed the MedusaDock simulations; performed statistical and chemical calculations; and created the figures. JW wrote the protein pocket extraction software and aided with the MedusaDock simulations. All three authors contributed to discussions regarding architecture exploration and design, the cost function, and data analysis and interpretation.

## REFERENCES

1. DiMasi, J. A., Grabowski, H. G. & Hansen, R. W. Innovation in the pharmaceutical industry: New estimates of R&D costs. J. Health Econ. 47, (2016).

2. Eder, J. & Herrling, P. L. Trends in modern drug discovery. in Handbook of Experimental Pharmacology vol. 232 (2016).

3. Goodnow, R. A., Dumelin, C. E. & Keefe, A. D. DNA-encoded chemistry: Enabling the deeper sampling of chemical space. Nature Reviews Drug Discovery vol. 16 (2017).

4. Waring, M. J. et al. An analysis of the attrition of drug candidates from four major pharmaceutical companies. Nature Reviews Drug Discovery vol. 14 (2015).

5. Hutchinson, L. & Kirk, R. High drug attrition rates - Where are we going wrong? Nature Reviews Clinical Oncology vol. 8 (2011).

6. Trott, O. & Olson, A. J. AutoDock Vina: Improving the speed and accuracy of docking with a new scoring function, efficient optimization, and multithreading. J. Comput. Chem. (2009) doi:10.1002/jcc.21334.

7. Allen, W. J. et al. DOCK 6: Impact of new features and current docking performance. J. Comput. Chem. 36, (2015).

8. How long does it take to screen 10,000 compounds with Glide? Schrödinger LLC. https://www.schrodinger.com/kb/1012 (2020).

9. Fan, M. et al. GPU-Accelerated Flexible Molecular Docking. J. Phys. Chem. B 125, (2021).

10. Bohacek, R. S., McMartin, C. & Guida, W. C. The art and practice of structure-based drug design: A molecular modeling perspective. Medicinal Research Reviews vol. 16 (1996).

11. Jumper, J. et al. High Accuracy Protein Structure Prediction Using Deep Learning. in Fourteenth Critical Assessment of Techniques for Protein Structure Prediction (Abstract Book) (2020).

12. Ding, F., Yin, S. & Dokholyan, N. V. Rapid flexible docking using a stochastic rotamer library of ligands. J. Chem. Inf. Model. 50, (2010).

13. Wang, J. & Dokholyan, N. V. MedusaDock 2.0: Efficient and Accurate Protein-Ligand Docking with Constraints. J. Chem. Inf. Model. 59, (2019).

14. Forli, S. et al. Computational protein-ligand docking and virtual drug screening with the AutoDock suite. Nat. Protoc. 11, (2016).

15. Goodsell, D. S., Sanner, M. F., Olson, A. J. & Forli, S. The AutoDock suite at 30. Protein Sci. 30, (2021).

16. Friesner, R. A. et al. Glide: A New Approach for Rapid, Accurate Docking and Scoring. 1. Method and Assessment of Docking Accuracy. J. Med. Chem. 47, (2004).

17. Evangelista, W. et al. Ensemble-based docking: From hit discovery to metabolism and toxicity predictions. Bioorganic Med. Chem. 24, (2016).

18. What is ensemble docking and how can I use it? Schrödinger LLC. https://www.schrodinger.com/kb/28 (2016).

19. Rocktäschel, T. & Riedel, S. End-to-end differentiable proving. in Advances in Neural Information Processing Systems vols 2017-December (2017).

20. Feng, Y., Zheng, Z., Liu, Q., Greenspan, M. & Zhu, X. Exploring end-to-end differentiable natural logic modeling. arXiv (2020) doi:10.18653/v1/2020.coling-main.101.

21. Basu, J. K., Bhattacharyya, D. & Kim, T. Use of Artificial Neural Network in Pattern Recognition. Int. J. Softw. Eng. its Appl. 4, (2010).

22. He, X., Zhu, T., Wang, X., Liu, J. & Zhang, J. Z. H. Fragment quantum mechanical calculation of proteins and its applications. Acc. Chem. Res. 47, (2014).

23. Francoeur, P. G. et al. Three-dimensional convolutional neural networks and a crossdocked data set for structure-based drug design. J. Chem. Inf. Model. 60, (2020).

24. Gentile, F. et al. Deep Docking: A Deep Learning Platform for Augmentation of Structure Based Drug Discovery. ACS Cent. Sci. 6, (2020).

25. Jiménez, J., Škalič, M., Martínez-Rosell, G. & De Fabritiis, G. KDEEP: Protein-Ligand Absolute Binding Affinity Prediction via 3D-Convolutional Neural Networks. J. Chem. Inf. Model. 58, (2018).

26. Cang, Z., Mu, L. & Wei, G. W. Representability of algebraic topology for biomolecules in machine learning based scoring and virtual screening. PLoS Comput. Biol. 14, (2018).

27. Yin, S., Biedermannova, L., Vondrasek, J. & Dokholyan, N. V. MedusaScore: An accurate force field-based scoring function for virtual drug screening. J. Chem. Inf. Model. 48, (2008).

28. Finzi, M., Stanton, S., Izmailov, P. & Wilson, A. G. Generalizing convolutional neural networks for equivariance to lie groups on arbitrary continuous data. arXiv (2020).

29. Wu, Z. et al. A Comprehensive Survey on Graph Neural Networks. IEEE Trans. Neural Networks Learn. Syst. 32, (2021).

30. Goodfellow, I. J. et al. Generative adversarial nets. in Advances in Neural Information Processing Systems vol. 3 (2014).

31. de Cao, N. & Kipf, T. MolGAN: An implicit generative model for small molecular graphs. arXiv (2018).

32. Schoenholz, S. S. & Cubuk, E. D. END-TO-END DIFFERENTIABLE, HARDWARE ACCELERATED, MOLECULAR DYNAMICS IN PURE PYTHON. arXiv (2019).

33. Wang, R., Fang, X., Lu, Y. & Wang, S. The PDBbind database: Collection of binding affinities for protein-ligand complexes with known three-dimensional structures. J. Med. Chem. 47, (2004).

34. Rdkit: Open-source chemoinformatics.

35. Irwin, J. J. & Shoichet, B. K. ZINC - A free database of commercially available compounds for virtual screening. J. Chem. Inf. Model. 45, (2005).

36. Berman, H. M. et al. The Protein Data Bank. Nucleic Acids Research vol. 28 (2000).

37. Miyato, T., Kataoka, T., Koyama, M. & Yoshida, Y. Spectral normalization for generative adversarial networks. in 6th International Conference on Learning Representations, ICLR 2018 - Conference Track Proceedings (2018).

38. Srivastava, N., Hinton, G., Krizhevsky, A., Sutskever, I. & Salakhutdinov, R. Dropout: A simple way to prevent neural networks from overfitting. J. Mach. Learn. Res. 15, (2014).

39. Maas, A. L., Hannun, A. Y. & Ng, A. Y. Rectifier nonlinearities improve neural network acoustic models. in in ICML Workshop on Deep Learning for Audio, Speech and Language Processing (2013).

40. He, K., Zhang, X., Ren, S. & Sun, J. Deep residual learning for image recognition. in Proceedings of the IEEE Computer Society Conference on Computer Vision and Pattern Recognition vols 2016-December (2016).

41. Abadi, M. et al. TensorFlow: A system for large-scale machine learning. in Proceedings of the 12th USENIX Symposium on Operating Systems Design and Implementation, OSDI 2016 (2016).

42. Kingma, D. P. & Ba, J. L. Adam: A method for stochastic optimization. in 3rd International Conference on Learning Representations, ICLR 2015 - Conference Track Proceedings (2015).

43. Szegedy, C. et al. Going deeper with convolutions. in Proceedings of the IEEE Computer Society Conference on Computer Vision and Pattern Recognition vols 07-12-June-2015 (2015).

44. Lipinski, C. A., Lombardo, F., Dominy, B. W. & Feeney, P. J. Experimental and computational approaches to estimate solubility and permeability in drug discovery and development settings. Advanced Drug Delivery Reviews vol. 64 (2012).

45. Bickerton, G. R., Paolini, G. V., Besnard, J., Muresan, S. & Hopkins, A. L. Quantifying the chemical beauty of drugs. Nat. Chem. 4, (2012).

46. Virtanen, P. et al. SciPy 1.0: fundamental algorithms for scientific computing in Python. Nat. Methods 17, (2020).

47. Vallat, R. Pingouin: statistics in Python. J. Open Source Softw. 3, (2018).

48. The PyMOL Molecular Graphics System, Version 2.4 Schrödinger, LLC.

