## Supplementary Information for NeuralDock for "NeuralDock: Rapid and conformation-agnostic docking of small molecules"

MedusaDock 2.0 augments the MedusaScore force field to calculate interaction energy terms^1^. We list the 13 interaction energy terms our models were trained on.

Table 1: Description of the 13 energies output by MedusaDock 2.0.

| **Energy term** | **Description** |
| --- | --- |
| E_total_ | The total binding energy of the protein and the ligand, computed as a difference from the energies of protein and ligand separately. |
| E_without VDWR_ | The total binding energy, but without repulsive van der Waals forces between the ligand and receptor. This energy has been seen to be more predictive of binding affinities. |
| E_VDWA_ | The attractive van der Waals forces. |
| E_VDWR_ | The repulsive van der Waals forces. |
| E_solv_ | The energy of solvation, using an EEF1 pairwise implicit solvent. |
| E_SB_ | The energies of bonds between side chain and backbone atoms. |
| E_HBBB_* | Hydrogen bonding in the backbone atoms. 0 for most structures. |
| E_HBSB_ | Hydrogen bonds between backbone and side chain atoms. |
| E_HBSS_ | Hydrogen bonds between side chain atoms. |
| E_FYAA_* | The backbone dihedral energy. 0 for most structures. |
| E_FYCHI_* | The side chain dihedral energy. 0 for most structures. |
| E_constraints_* | An energy cost to impose constraints. 0 for all structures. |
| E_AAREF_* | Additional terms for specific amino acid interactions, such as proline, glycine, and cysteine. 0 for most structures. |

Table 2: Correlation coefficients for each of the predicted energy statistics for the 45 million parameter network. The minimum E_without VDWR_ is the scatter data shown in Figure 1b.

| **Correlation coefficients** | | | | | | | | |
| --- | --- | --- | --- | --- | --- | --- | --- | --- |
|  | **E_total_** | **E_without VDWR_** | **E_VDWA_** | **E_VDWR_** | **E_solv_** | **E_SB_** | **E_HBSB_** | **E_HBSS_** |
| mean | 0.477046 | 0.775044 | 0.9047 | 0.358243 | 0.788617 | 0.532118 | 0.41008 | 0.707306 |
| std | 0.06842 | 0.637904 | 0.688438 | 0.071204 | 0.599799 | 0.150192 | 0.389633 | 0.500484 |
| median | 0.670155 | 0.769709 | 0.897751 | 0.696402 | 0.78447 | 0.574416 | 0.358121 | 0.676497 |
| min | 0.78836 | **0.837795** | 0.928523 | 0.506693 | 0.728322 | 0.520312 | 0.503994 | 0.661118 |
| max | 0.154187 | 0.689073 | 0.821365 | 0.153536 | 0.77941 | 0.578034 | 0 | 0.02498 |
| skew | 0.246509 | 0.2133 | 0.253677 | 0.143483 | 0.26284 | 0.165613 | 0.168141 | 0.29438 |
| kurtosis | 0.185091 | 0.235944 | 0.303604 | 0.096907 | 0.447759 | 0.270723 | 0.136571 | 0.148129 |
